# HCV infection induces ubiquitin-dependent degradation of LATS1, inactivating the Hippo pathway and upregulating transcription of the CYR61 and CTGF genes

**DOI:** 10.1101/2025.04.08.647823

**Authors:** Maria Alethea Septianastiti, Chieko Matsui, Zihan Xu, Fransisca Puspitasari, Dewa Nyoman Murti Adyaksa, Lin Deng, Takayuki Abe, Ikuo Shoji

## Abstract

Hepatitis C virus (HCV) is often associated with chronic liver diseases and significant alterations in host cellular signaling. However, the molecular mechanisms underlying HCV- related liver pathogenesis remain to be elucidated. The Hippo signaling pathway, a key regulator of cell proliferation and survival, plays a critical role in maintaining liver homeostasis. Here we investigated the role of the Hippo pathway in HCV-related pathogenesis. We demonstrated that HCV infection induces degradation of LATS1, a key regulator of the Hippo pathway. Degradation of LATS1 protein was restored by a proteasomal inhibitor, but not a lysosome inhibitor, indicating that HCV promotes proteasomal degradation of LATS1 protein. HCV-induced degradation of LATS1 protein was suppressed in si-Itch-transfected Huh-7.5 cells. These results suggest that Itch ubiquitin ligase is involved in ubiquitin-dependent degradation of LATS1 protein. Cell fractionation assays and immunofluorescence staining revealed that HCV infection promoted nuclear translocation of YAP1 protein, suggesting that HCV infection suppresses the Hippo pathway. Furthermore, the transcription of YAP1 target genes, CYR61 and CTGF, that are involved in tissue remodeling and proliferation, was upregulated in HCV-infected Huh-7.5 cells and in HCV-infected patients. Taken together, we propose that HCV promotes the ubiquitin-dependent proteasomal degradation of LATS1 protein, leading to suppression of the Hippo pathway, thereby upregulating transcription of CYR61 and CTGF genes.

**DATA SUMMARY:** All data are presented in the main figures. Raw sequencing data, microscopy images, materials, and sequence information are available upon request. Correspondence and requests for materials should be addressed to Professor Ikuo Shoji. The data that support the findings of this study is available at bioRxiv (https://www.biorxiv.org/).

**IMPACT STATEMENT:** We demonstrate evidence suggesting that HCV infection promotes the Itch-mediated ubiquitin-dependent degradation of LATS1 protein, a key factor for the Hippo pathway. HCV-induced ubiquitin-dependent degradation of LATS1 protein promotes inactivation of the Hippo pathway and nuclear translocation of YAP1 protein, thereby upregulating transcription of CYR61 and CTGF genes. We propose a novel molecular mechanism in which HCV infection promotes degradation of LATS1 protein, leading to inactivation of the Hippo pathway. Understanding HCV-induced inactivation of the Hippo pathway may lead to developing new strategies for preventing or treating HCV-related pathogenesis.

## INTRODUCTION

Hepatitis C virus (HCV) infection remains a global health burden, affecting an estimated 50 million people worldwide as of 2022 (1). HCV infection often causes chronic liver diseases, ranging from mild illness to severe complications, including liver cirrhosis and hepatocellular carcinoma (HCC). Although highly effective direct-acting antivirals (DAAs) are available for HCV treatment, the risk of HCC persists even after successful viral clearance (2). Notably, HCC can occur following a sustained virological response, at an annual incidence of approximately 1%, surpassing the cancerous conditions in other organs (3). HCV is a positive-sense, single-stranded RNA virus classified in the *Flaviviridae* family, *Hepacivirus* genus. The 9.6-kb HCV RNA genome encodes a single polyprotein of approximately 3,010 amino acids, which is cleaved by viral proteases and host signalases into structural proteins (Core, E1, and E2) and nonstructural proteins (p7, NS2, NS3, NS4A, NS5A, and NS5B) (4).

Understanding HCV-induced modulation of host cellular signaling pathway is crucial for elucidating mechanisms that drive liver disease progression. The Hippo pathway, a key regulator of cell proliferation, apoptosis, and organ size control, plays an essential role in tissue homeostasis. Dysregulation of the Hippo pathway is implicated in the development of several cancers, including liver cancer (5–7). The Hippo pathway involves a series of proteins that form a kinase cascade, including MST1/2, LATS1/2, the adaptor protein SAV1, MOB1A/B, and the transcription coactivators YAP1 and TEAD transcription factors. Activation of the Hippo pathway involves the phosphorylation of the Serine/Threonine kinase LATS1, which in turn phosphorylates YAP1 at Ser127 and promotes the binding of phosphorylated YAP1 to 14-3-3 proteins (8, 9). The association of 14-3-3 proteins with phosphorylated YAP1 prevents it from entering the nucleus. Conversely, when the Hippo pathway is inactivated, unphosphorylated YAP1 translocates to the nucleus and induces the transcription of genes responsible for cell growth (10,11).

Several viruses have been shown to modulate the Hippo pathway (12–15). HCV E2 activates the Hippo pathway by interacting with CD81, while HCV NS4B suppresses the Hippo pathway via Scribble, promoting PI3K/AKT signaling and epithelial-mesenchymal transition (EMT) (16,17). Similarly, Kaposi sarcoma-associated herpesvirus (KSHV) vGPCR inhibits LATS1/2, enabling YAP1 nuclear translocation. Human papillomavirus 16 E6 alters YAP1 localization through interaction with PDZ domains (13,15).

We previously reported that HCV infection activates the ROS/JNK signaling pathway which in turn activates Itch, a HECT-type E3 ubiquitin ligase. Activated Itch promotes polyubiquitylation of VPS4A, resulting in efficient HCV particle release (18). Notably, Itch has been shown to promote the proteasomal degradation of LATS1, a critical component of the Hippo pathway (19,20). In this study, we investigate a role of the Hippo pathway in HCV-related pathogenesis. We demonstrate that Itch plays an important role in HCV-induced ubiquitin-dependent degradation of LATS1, leading to Hippo pathway suppression and thereby contributing to HCV pathogenesis.

## METHODS

### Cell culture and viruses

Human hepatoma cell line Huh-7.5 cells were generously provided by Dr. Charles M. Rice (The Rockefeller University, New York, NY). The cells were cultured in Dulbecco’s Modified Eagle’s Medium (DMEM) (high glucose) containing L-glutamine and phenol red (044-29765; Fuji Film Wako Pure Chemical Industries, Osaka, Japan) supplemented with 50 IU/ml penicillin, 50 µg/ml streptomycin (15-140-122; Gibco, Grand Island, NY), 10% heat-inactivated fetal bovine serum (FBS) (S1760-500; Biowest, Nuaillé, France), and 0.1 mM non-essential amino acids (Invitrogen, Carlsbad, CA). Cells were cultured at 37°C in a 5% CO incubator. Cells were washed with PBS (-) solution (05913; Nissui Pharmaceutical Co., Ltd., Tokyo, Japan) and transfected with plasmid DNA using FuGENE 6 transfection reagents (E269A; Promega, Madison, WI) following the manufacturer’s instructions. The pFL-J6/JFH1 plasmid, encoding the entire viral genome of the chimeric HCV-2a strain J6/JFH1, was also kindly provided by Dr. Charles M. Rice. HCV genome RNA was synthesized *in vitro* using pFL-J6/JFH1 as a template and subsequently transfected into Huh-7.5 cells via electroporation. The virus produced in the culture supernatant was then used for infection experiments (21, 22).

The Huh-7 cells stably harboring HCV genotype 1b (RCYM1) were described previously (23)

### Expression plasmids

The cDNA fragment of LATS1 was amplified by PCR using the following specific primers: forward 5’-TCGAGCTCAGCGGCCATGAAGAGGAGTGAAAAGCCAG-3’ and reverse 5’-TCGAGCTCAGCGGCCATGAAGAGGAGTGAAAAGCCAG-3’. The amplified PCR product was purified and inserted into the NotI site of the pCAG-FLAG plasmid using the In-Fusion HD Cloning Kit (Clontech, Mountain View, CA). The sequences of the inserts were verified by sequencing (Eurofins Genomics, Tokyo, Japan).

To construct the pCAG-Myc-LATS1 expression plasmid, pCAG-FLAG-LATS1 was used as a template and the cDNA fragment of LATS1 was amplified using the following specific primers: forward 5’-AAGGTACCATGGAGCAAAAGCTCATTTCTGAAGAGGACCTGATGAAGAGGAGTGAAAAGCCA-3’ and reverse 5’-TTAGATCTTCAAACATATACTAGATCGCGATT-3’. The amplified PCR product was purified and inserted into the KpnI and BglII sites of the pCAG-MCS2 vector using the Ligation-Convenience Kit (Nippon Gene, Tokyo, Japan).

The expression plasmids pCAG-FLAG-Itch and pCAG-FLAG-WWP1 were described previously (24). The C890A point mutant of WWP1 was generated by overlap extension PCR using pCAG-FLAG-WWP1 as a template and the following specific primers: forward 5’-AGAAGCCATACAGCTTTTAATCGCTTG-3’ and reverse 5’-CAAGCGATTAAAAGCTGTATGGCTTCT-3’. The sequences of the inserts were verified by sequencing (Eurofins Genomics).

N-terminal HA-tagged ubiquitin (Ub) expression plasmid was purchased from Addgene (Watertown, MA).

### Antibodies and reagents

The mouse monoclonal antibodies (MAbs) used in this study were anti-FLAG (M2) MAb (F3165, Sigma-Aldrich), anti-β-actin (A5441; Sigma-Aldrich) anti-c-Myc (9E10) Mab (sc-40; Santa Cruz Biotechnology, Dallas, TX), anti-Itch Mab (611198, BD Transduction Laboratories, San Jose, CA), anti-Histone H3 (1G1) MAb (sc-517576; Santa Cruz Biotechnology), and anti-YAP (63.7) MAb (sc-101199; Santa Cruz Biotechnology). The rabbit monoclonal antibody used in this study was anti-LATS1 (C66B5) MAb (3477; Cell Signaling Technology). The rabbit polyclonal antibodies (PAbs) used in this study included anti-HA PAb (H-6908; Sigma-Aldrich), anti-phospho-LATS1 (Ser909) PAb (9157; Cell Signaling Technology), anti-LAMP2A PAb (ab18528; Abcam, Cambridge, UK), anti-YAP PAb (4912; Cell Signaling Technology), anti-phospho-YAP (Ser127) (4911; Cell Signaling Technology), anti-NS5A PAb (2914-1; a kind gift from T. Wakita, National Institute of Infectious Diseases, Tokyo, Japan), and anti-IκBα PAb (9242; Cell Signaling Technology). Horseradish peroxidase (HRP)-conjugated anti-mouse IgG (7076S; Cell Signaling Technology) and HRP-conjugated anti-rabbit IgG (7074S; Cell Signaling Technology) were used as secondary antibodies. Clasto-lactacystin β-lactone (031-18201; Wako) was used as a proteasome inhibitor. Ammonium chloride (NH_4_Cl) was purchased from Fuji Film Wako Pure Chemical Industries.

### Immunoblot analysis

Immunoblot analysis was conducted as previously described (25). Cell lysates were separated using 8% or 15% sodium dodecyl sulfate−polyacrylamide gel electrophoresis (SDS-PAGE) and transferred onto a 0.45 µm Immobilon-P polyvinylidene difluoride (PVDF) membrane (Millipore, Billerica, MA). The membranes were incubated with a primary antibody, followed by an HRP-conjugated secondary antibody. Positive bands were detected using Amersham enhanced chemiluminescence (ECL) western blotting detection reagents (Cytiva, RPN2106; Sigma-Aldrich). Band intensities were quantified using ImageJ software (version 1.54h).

### Cell-based ubiquitylation assay and immunoprecipitation

Cell-based ubiquitylation assays and immunoprecipitation experiments were performed as previously described (24). Cultured cells were lysed in a lysis buffer containing 50 mM HEPES (pH 7.5), 150 mM NaCl, 1 mM EDTA, 1% NP-40, 1 mM DTT, 1 mM PMSF, 10 mM pyrophosphate, 10 mM glycerophosphate, 50 mM NaF, 1.5 mM Na[VO[, and a complete™, EDTA-free protease inhibitor cocktail (Roche Molecular Biochemicals, Mannheim, Germany). After sonication, the lysates were immunoprecipitated using either anti-FLAG M2 affinity gel (A2220; Sigma-Aldrich) or Protein A-Sepharose 4 Fast Flow (GE17-5280-04; GE Healthcare, Buckinghamshire, UK) pre-incubated with the appropriate antibodies. Immunoprecipitation was carried out at 4°C for 4 h or overnight. The immunoprecipitates were washed five times with the lysis buffer and analyzed by immunoblotting using specific antibodies, including anti-HA for ubiquitylation assays.

### Cell fractionation assay

Cytoplasmic and nuclear extracts from HCV-infected and mock-infected Huh-7.5 cells were separated using NE-PER™ Nuclear and Cytoplasmic Extraction Reagents (78835; Thermo Fisher Scientific) following the manufacturer’s instructions. The resulting lysates were then analyzed by immunoblotting.

### CHX chase experiment

To determine the half-life of LATS1 protein, cells were treated with 50 µg/mL cycloheximide (CHX). Cells designated as the zero-time point were harvested immediately after CHX treatment. For subsequent time points, cells were incubated CHX-containing medium at 37°C for 6, 12, 18, and 24 hours, as indicated.

### RNA extraction and RT-qPCR

Total RNA was extracted using the ReliaPrep RNA Cell Miniprep System (Promega) following the manufacturer’s protocol. The GoScript Reverse Transcription System (Promega) was used for cDNA synthesis. RT-qPCR was performed on a StepOnePlus Real-Time PCR System (Applied Biosystems, Foster City, CA) using TB Green Premix Ex Taq II (TaKaRa Bio, Shiga, Japan) with SYBR Green chemistry. The primer sequences used were as follows: LATS1 (forward: 5’-CTCTGCACTGGCTTCAGATG-3’, reverse: 5’-TCCGCTCTAATGGCTTCAGT-3’); YAP1 (forward: 5’-CCCGACAGGCCAGTACTGAT-3’, reverse: 5’-CAGAGAAGCTGGAGAGGAATGAG-3’); CYR61 (forward: 5’-GGTCAAAGTTACCGGGCAGT-3’, reverse: 5’-GGAGGCATCGAATCCCAGC-3’); and CTGF (forward: 5’-CCTGGTCCAGACCACAGAGT-3’, reverse: 5’-ATGTCTTCATGCTGGTGCAG-3’).

Expression of the human GAPDH gene was measured as an internal control using the primers (forward: 5’-GCCATCAATGACCCCTTCATT-3’, reverse: 5’-TCTCGCTCCTGGAAGATGG-3’.

### RNA interference and stable knockdown cells

The generation of shControl and shLAMP2A cells was described previously (26). HCV-infected and uninfected Huh-7.5 cells were transfected with either Itch siRNA (SI00141085; Qiagen,). WWP1 siRNA (JP00049582-001; Sigma-Aldrich), YAP siRNA (WD11034049-001; Sigma-Aldrich), or negative-control scrambled siRNA (1027281; Qiagen) using Lipofectamine RNAiMAX transfection reagent (Life Technologies, Carlsbad, CA) according to the manufacturer’s instructions. The siRNA sequences were as follows: siWWP1, 5’-CUUCUACGAUCAUCAACUCTT-3’; siYAP1, 5’-TTCTTTATCTAGCTTGGTGGC-3’. All siRNAs were transfected once.

### Immunofluorescence staining

Huh-7.5 cells cultured on glass coverslips were fixed with 4% paraformaldehyde at room temperature (RT) for 15 min. After being washed with phosphate-buffered saline (PBS) (05913; Nissui Pharmaceutical Co., Ltd.), the cells were permeabilized with PBS containing 0.1% Triton X-100 for 15 min at RT. To block nonspecific binding, the cells were incubated in PBS containing 1% bovine serum albumin (BSA) (01859-47; Nacalai Tesque) for 60 min. The cells were then incubated with 1% BSA in PBS containing mouse anti-YAP monoclonal antibody and rabbit anti-NS5A polyclonal antibody at RT for 60 min. Afterward, the cells were washed three times with PBS and further incubated with 1% BSA in PBS containing Alexa Fluor™ 594-conjugated anti-mouse IgG (A11005; Invitrogen) and Alexa Fluor™ 488-conjugated anti-rabbit IgG (A11008; Invitrogen) at RT for 60 min. Finally, the cells were washed four times with PBS, mounted on glass slides, and examined using a confocal microscope (LSM 700, Carl Zeiss, Oberkochen, Germany).

### Transcriptomic data analysis

To evaluate the expression of CYR61 and CTGF mRNAs in HCV-infected liver tissue, transcriptomic data were obtained from the GEO database (accession number GSE84346). The dataset included samples from HCV-infected patients (n = 38) and healthy controls (n = 6). Normality of gene expression data was assessed using the Kolmogorov-Smirnov and Shapiro-Wilk tests. As the data were not normally distributed, group comparisons were performed using the non-parametric Mann-Whitney U test. A p-value <0.05 was considered statistically significant.

### Statistical analysis

For RT-qPCR data, results were expressed as means ± standard error of the mean (SEM). Statistical significance was evaluated using Student’s t-test and was defined as a P value of <0.05.

## RESULTS

### HCV promotes proteasomal degradation of LATS1 protein

To investigate a role of the Hippo signaling pathway in HCV-related pathogenesis, we examined LATS1 protein levels in HCV J6/JFH1-infected Huh-7.5 cells. Immunoblot analysis revealed a reduction in LATS1 protein levels in HCV-infected cells at both 2 and 4 days post-infection (Fig. 1A, first panel, lanes 2 and 4). However, reverse-transcription quantitative polymerase chain reaction (RT-qPCR) analysis showed increased LATS1 mRNA levels (Fig. 1B), indicating that HCV-induced reduction of LATS1 protein levels is not due to suppression of LATS1 mRNA expression.

**Figure 1.**
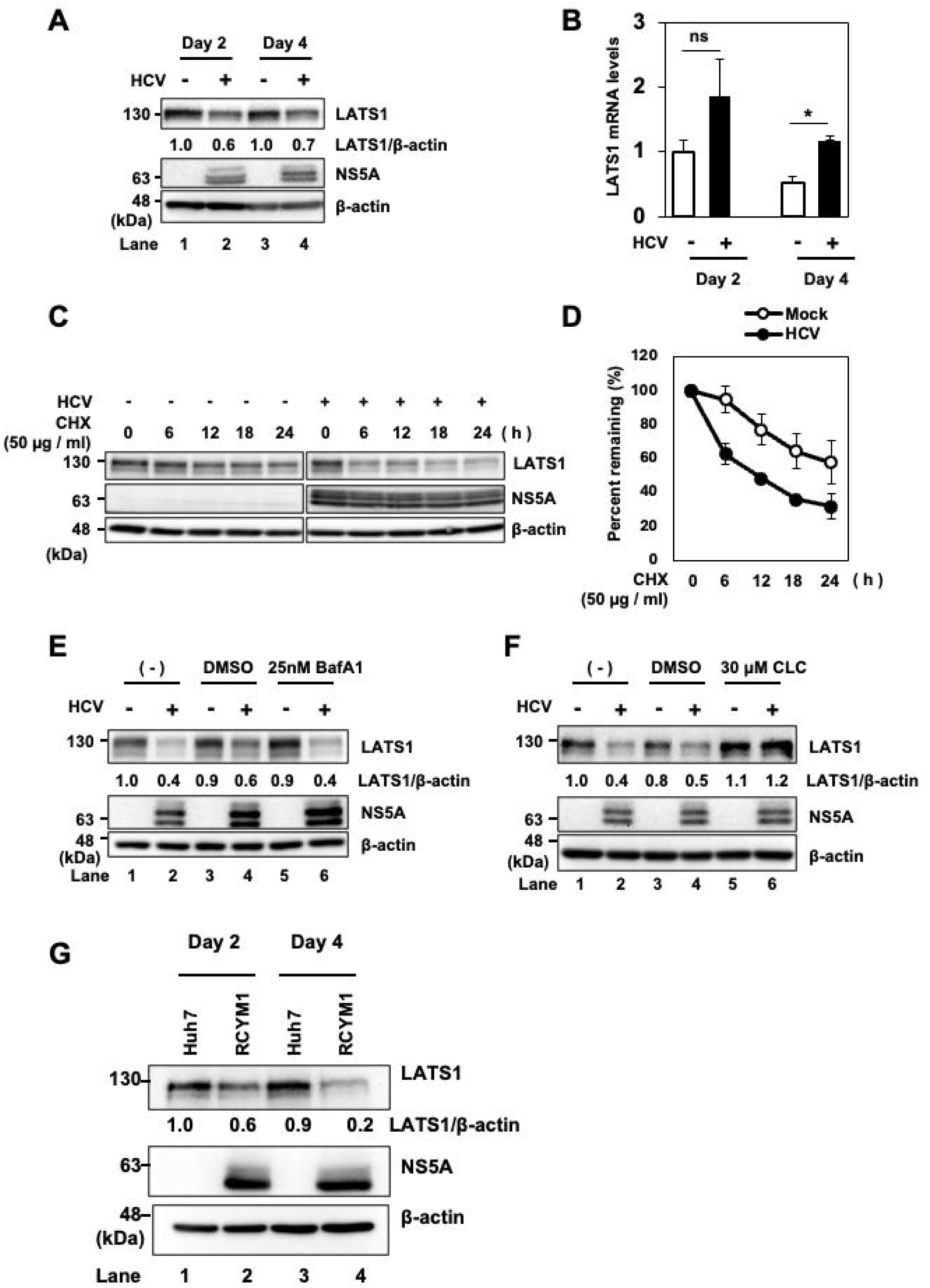
HCV induces proteasomal degradation of LATS1 protein. (A) Huh-7.5 cells were infected with HCV J6/JFH1 at a multiplicity of infection (MOI) of 2. Cells were cultured and harvested at 2 and 4 days post infection (dpi). Immunoblot analysis was performed using indicated antibodies. β-actin levels were used as a loading control. (B) Huh-7.5 cells were infected with HCV J6/JFH1 at an MOI of 2. Cells were cultured and harvested at 2 and 4 days dpi. Subsequently, total cellular RNA was extracted, and LATS1 mRNA levels were quantified by RT-qPCR. To normalize the LATS1 mRNA levels, GAPDH mRNA levels were used as an internal control. The value for day 2 in the mock-infected group was arbitrarily expressed as 1.0. Data are presented as means ± SEM from three independent experiments that yielded similar results; p-value < 0.05 (*) was significant, compared with the controls. (C) HCV-infected and mock-infected control cells were treated with 50 µg/mL cycloheximide (CHX) for 0, 6, 12, 18, and 24 hours. Immunoblot analysis was performed using indicated antibodies, and β-actin levels served as a loading control. (D) Specific signals were quantified by densitometry. The percentages of remaining LATS1 and β-actin at each time point were normalized to their respective basal level. Closed circles represent HCV-infected cells and open circles represent mock-infected control cells. Data shown are representative of three independent experiments that yielded similar results. (E) Huh-7.5 cells were infected with HCV J6/JFH1 at an MOI of 2, followed by treatment with Bafilomycin A1 for 12 hours. Immunoblotting was performed with indicated antibodies and β-actin levels served as a loading control. (F) Huh-7.5 cells were infected with HCV J6/JFH1 at an MOI of 2. Cells were treated with 30µM Clasto-Lactacystin β-Lactone for 12 hours, harvested and analyzed by immunoblotting using the indicated antibodies. β-actin served as a loading control. The immunoblots are representative of three independent experiments that yielded similar results. (G) Huh-7 cells stably harboring the HCV genotype genome-length replicon (RCYM1) and the parental Huh-7 cells were seeded and cultured for 2 or 4 days. Cells were harvested and analyzed by immunoblotting using the indicated antibodies.

To determine whether HCV-induced reduction of LATS1 protein is protein degrdataion, we performed CHX-chase analyses. The CHX-chase analyses revealed that the half-life of LATS1 protein was significantly reduced upon HCV infection (Fig. 1C, first panel; Fig. 1D).

To further elucidate a mechanism underlying HCV-induced reduction of LATS1 protein, we treated HCV-infected Huh-7.5 cells with the lysosomal inhibitor bafilomycin A1. The LATS1 protein levels in HCV-infected cells were not restored by treatment with bafilomycin A1 (Fig. 1E, first panel, lane 6), suggesting that HCV-induced LATS1 protein reduction is not due to the lysosomal degradation.

To determine whether HCV-induced LATS1 protein reduction is due to proteasomal degradation, we treated the cells with the proteasome inhibitor clasto-lactacystin β-lactone. Treatment of the cells with clasto-lactacystin β-lactone restored LATS1 protein levels in HCV-infected cells (Fig. 1F, first panel, lane 6), indicating that HCV-induced LATS1 protein reduction is due to proteasomal degradation. Collectively, these results indicate that HCV promotes proteasomal degradation of LATS1 protein.

To determine whether HCV-induced LATS1 protein reduction is conserved among different HCV genotypes, we utilized Huh-7 cells harboring the HCV genotype 1b genome-length replicon (RCYM1). Immunoblot analyses revealed that LATS1 protein levels were reduced compared to parental Huh-7 cells (Fig. 1G, first panel, lanes 2 and 4). These results indicate that LATS1 protein is reduced in HCV-1b replicating cells.

### E3 ligase Itch, but not WWP1, plays a crucial role in HCV-induced polyubiquitylation and degradation of LATS1

To determine whether HCV-induced LATS1 degradation depends specifically on Itch ubiquitin ligase, we generated an inactive mutant Itch by mutating cysteine 868 to alanine (pCAG-FLAG-Itch C868A), which abolishes E3 ligase activity. Huh-7.5 cells were co-transfected with pCAG-Myc-LATS1, pRK5-HA-Ub, and either pCAG-FLAG-Itch or pCAG-FLAG-Itch C868A. Immunoblot analysis revealed that LATS1 polyubiquitylation and degradation were increased upon co-transfection with pCAG-FLAG-Itch (Fig. 2A, first and second panel, lane 2) but not with the inactive mutant (Fig. 2A, first and second panel, lane 3). These results suggest that Itch mediates polyubiquitylation and degradation of LATS1.

**Figure 2.**
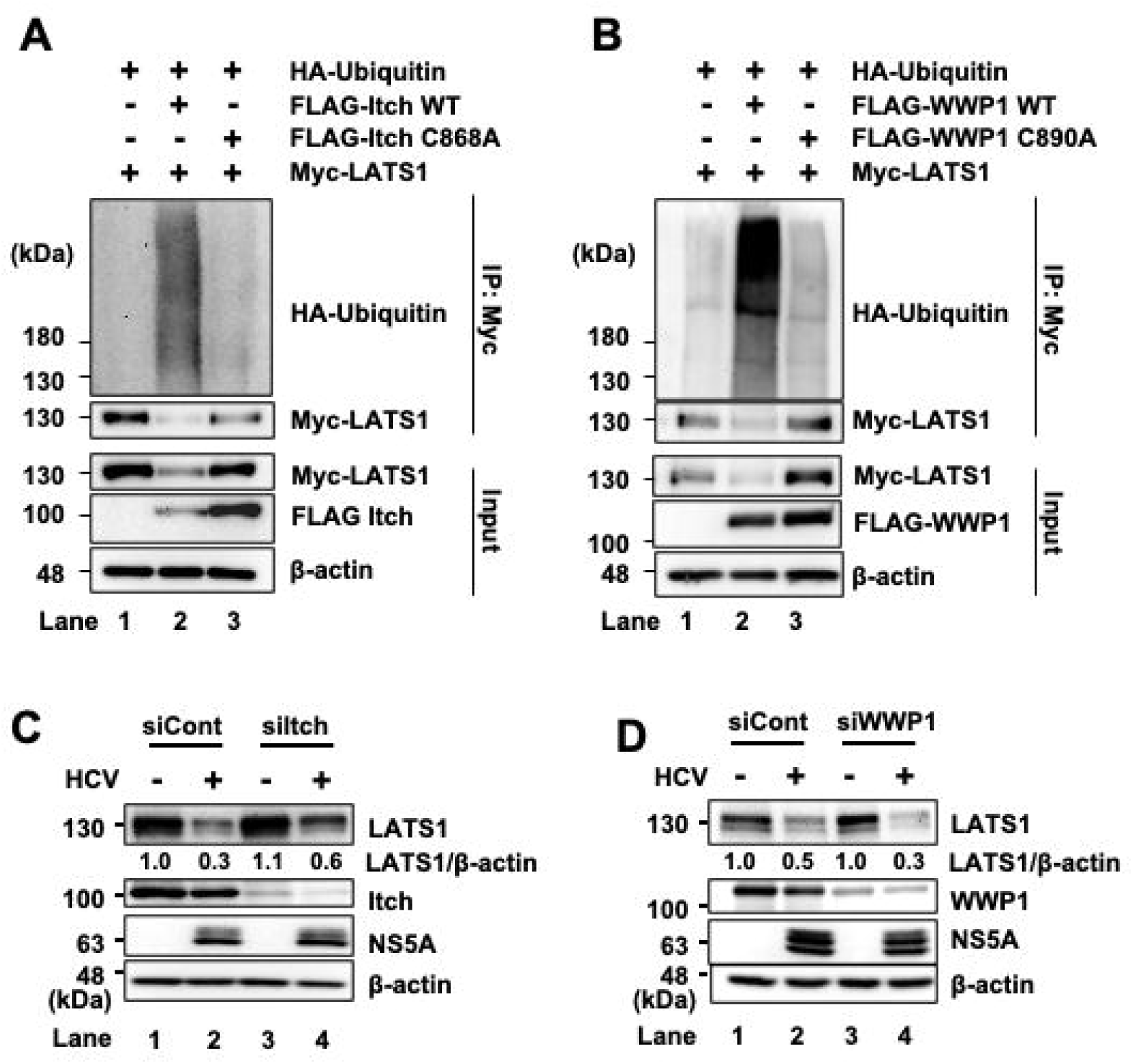
E3 ligase Itch, but not WWP1, plays a crucial role in HCV-induced polyubiquitylation and degradation of LATS1 protein. (A) Huh-7.5 cells were co-transfected with pCAG-Myc-LATS1, pRK5-HA-Ub, together with either pCAG-FLAG-Itch WT or pCAG-FLAG-Itch C868A. At 2 days after transfection, the cells were harvested. The cell lysates were immunoprecipitated with anti-c-Myc antibody, followed by immunoblotting using anti-HA rabbit pAb and anti-c-Myc mouse mAb. Input samples were also analyzed by immunoblotting using anti-c-Myc mouse mAb, and anti-FLAG mouse mAb. β-actin served as a loading control. (B) Huh-7.5 cells were co-transfected with pCAG-Myc-LATS1, pRK5-HA-Ubiquitin, together with either pCAG-FLAG-WWP1 WT or pCAG-FLAG-WWP1 C890A. At 2 days after transfection, cells were harvested. The cell lysates were immunoprecipitated with anti-c-Myc antibody, followed by immunoblotting with anti-HA rabbit pAb and anti-c-Myc mouse mAb. Input samples were also analyzed by immunoblotting using anti-c-Myc mouse mAb, and anti-FLAG mouse mAb. β-actin served as a loading control. (C) Huh-7.5 cells at 3 × 10^5^ cells in a 12-well plate were transfected with 48 pmol of either control siRNA or Itch-specific siRNA. After 24 hours, the cells were infected with HCV J6/JFH1 at an MOI of 2 and harvested at 2 days post-infection. The samples were analyzed by immunoblotting with specified antibodies. β-actin served as a loading control. (D) Huh-7.5 cells at 3 × 10^5^ cells in a 12-well plate were transfected with 48 pmol of either control siRNA or WWP1-specific siRNA. At 24 hours after siRNA-transfection, the cells were infected with HCV J6/JFH1 at an MOI of 2. Cells were harvested at 2 days post-infection and the samples were subjected to immunoblotting with the indicated antibodies. β-actin served as a loading control. The immunoblots are representative of three independent experiments that yielded similar results.

To determine whether other E3 ubiquitin ligases in the HECT family, such as WWP1, also play a role in the polyubiquitylation of LATS1, we performed a cell-based ubiquitylation assay using wild-type pCAG-FLAG−WWP1 and the inactive mutant pCAG-FLAG−WWP1 C890A. Immunoblot analysis showed that LATS1 polyubiquitylation and degradation were increased upon co-transfection with wild-type pCAG-FLAG-WWP1 (Fig. 2B, first and second panel, lane 2), while the inactive mutant abolishes this effect (Fig. 2B, first and second panel, lane 3).

To verify Itch’s role in HCV-induced LATS1 degradation, we performed siRNA-mediated knockdown of either Itch or WWP1 in HCV-infected cells. Immunoblot analysis demonstrated that knockdown of Itch restored LATS1 protein levels (Fig. 2C, first panel, lanes 2 and 4), whereas knockdown of WWP1 further decreased LATS1 protein levels (Fig. 2D, first panel, lanes 2 and 4). These findings suggest that Itch is responsible for HCV-induced ubiquitin-dependent degradation of LATS1.

### HCV infection promotes translocation of YAP1 from the cytoplasm to the nucleus

LATS1 is the protein kinase that phosphorylates YAP1 at Ser127 and activates the Hippo signaling pathway. Therefore, we hypothesized that HCV-induced degradation of LATS1 may suppress the Hippo pathway. To assess this hypothesis, we performed immunoblot analysis of components of the Hippo pathway, LATS1 and YAP1, in mock- and HCV J6/JFH1-infected Huh-7.5 cells. Immunoblot analysis revealed that HCV infection promoted LATS1 degradation (Fig. 3A, first panel, lane 2), accompanied by a reduction in phosphorylated LATS1 at Ser 909 (Fig. 3A, second panel, lane 2), suggesting that active LATS1 is decreased in HCV-infected cells. The decrease in pLATS1 at Ser 909 subsequently resulted in reduced phosphorylated YAP1 at Ser 127 (Fig. 3A, third panel, lane 2), whereas total YAP1 levels remained unchanged (Fig. 3A, fourth panel, lane 2). These results suggest that HCV infection reduced LATS1 and phosphorylated LATS1 at Ser 909, resulting in decreased phosphorylated YAP1 at Ser 127, thereby inactivating the Hippo pathway.

**Figure 3.**
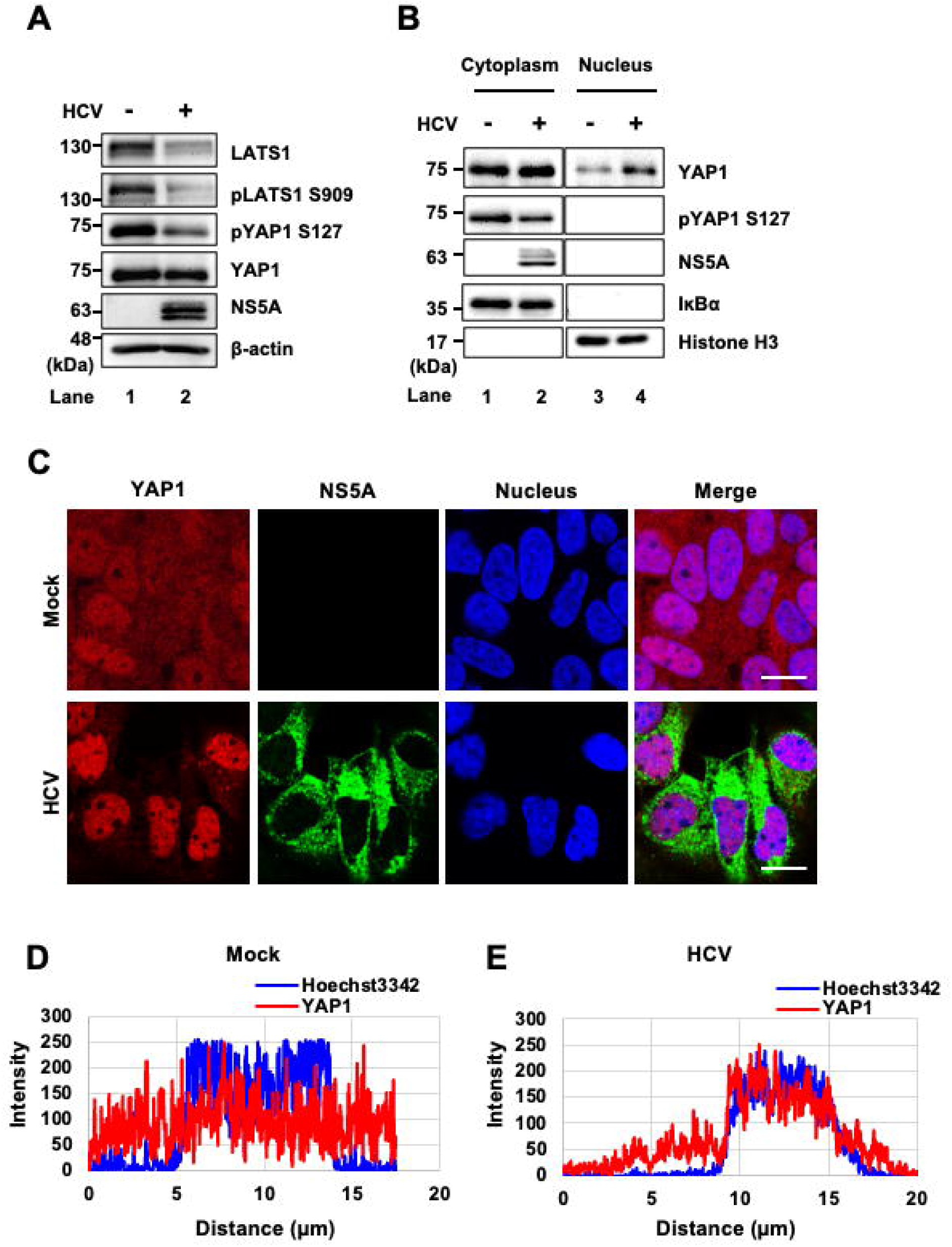
HCV infection promotes the translocation of YAP1 from the cytoplasm to the nucleus. (A) Huh-7.5 cells were infected with HCV J6/JFH1 at an MOI of 2. At 3 days after infection, the cells were harvested and analyzed by immunoblotting using the indicated antibodies targeting components of the Hippo pathway. β-actin served as a loading control. (B) Huh-7.5 cells were infected with HCVJ6/JFH1 at an MOI of 2. At 3 days after infection, the cells were harvested and subjected for cell fractionation analysis. IĸBα served as a loading control for the cytoplasmic fraction and Histone H3 served as a loading control for the nuclear fraction. The immunoblots are representative of three independent experiments that yielded similar results. (C) Huh-7.5 cells were plated and cultured for 12 hours and infected with HCV J6/JFH1 at an MOI of 2. Cells were stained with anti-YAP1 monoclonal antibody followed by Alexa Fluor 594-conjugated goat anti-mouse IgG (red) and anti-NS5A polyclonal antibody followed by Alexa Fluor 488-conjugated goat anti-rabbit IgG (green). Nuclei were counterstained with Hoechst 33342 (blue). Images were acquired using scanning laser confocal microscopy and processed with ImageJ software. Scale bar: 10µm. The immunofluorescence staining results are representative of three independent experiments. (D, E) A straight line was drawn from the cytoplasm through the nucleus of a representative cell in confocal immunofluorescence images from the mock (D) and HCV-infected (E) groups using ImageJ. The corresponding line scan profiles display the fluorescence intensity (gray value, arbitrary units) of YAP1 (red line) and Hoechst 33342 (blue line) along the measured distance. The degree of overlap between the YAP1 and Hoechst 33342 signals indicates the extent of YAP1 nuclear localization. Data shown are representative of three independent experiments.

To determine whether unphosphorylated YAP1 translocates to the nucleus, we performed a cell fractionation assay coupled with immunoblot analysis. The cell fractionation assay revealed an increased nuclear YAP1 in HCV-infected cells (Fig. 3B, first panel, lane 4) compared to mock-infected cells (Fig. 3B, first panel, lane 3). Furthermore, immunofluorescence staining revealed that YAP1 proteins were predominantly localized to the nucleus in HCV-infected Huh-7.5 cells (Fig. 3C, second panel). The line scan analysis showed that YAP1 exhibited cytoplasmic and nuclear distribution in mock-infected cells as shown by the fluorescence and the corresponding line scan profile (Fig. 3C, first panel; Fig. 3D). In contrast, the YAP1 intensity peak overlapped with the nuclear signal peak in HCV-infected cells, indicating YAP1 accumulation within the nucleus (Fig. 3C, second panel; Fig. 3E). These results suggest that HCV infection promotes LATS1 degradation, leading to reduced YAP1 phosphorylation and facilitating YAP1 nuclear translocation, thereby inactivating the Hippo pathway.

### mRNA levels of YAP1 target genes are upregulated upon HCV infection

To determine whether HCV-induced nuclear translocation of YAP1 affects downstream gene expression, we performed RT-qPCR to measure the mRNA levels of canonical YAP1 target genes, CYR61 and CTGF, in HCV-infected cells. RT-qPCR showed that HCV infection significantly increased the mRNA levels of CYR61 and CTGF (Fig. 4A and 4B).

**Figure 4.**
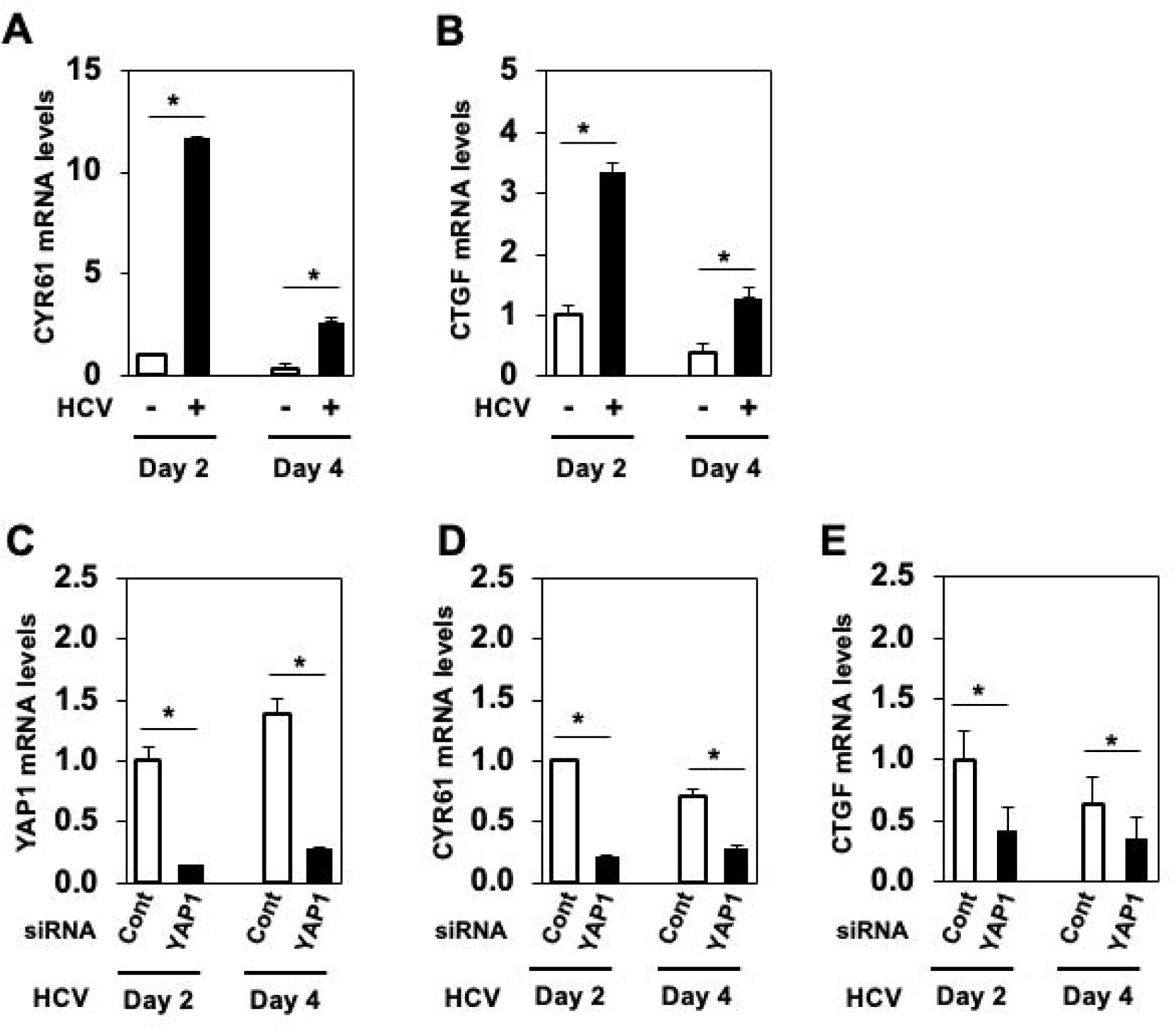
The mRNA levels of YAP1 target genes are significantly upregulated upon HCV infection. (A, B) Huh-7.5 cells were infected with HCV J6/JFH1 at an MOI of 2. Cells were cultured and harvested at 2 and 4 days post infection (dpi). Total cellular RNA was subsequently extracted, and the mRNA levels of CYR61 and CTGF were quantified by RT-qPCR. GAPDH mRNA levels were used for normalization. The mRNA level for day 2 in the mock-infected group was arbitrarily expressed as 1.0. Data are presented as means ± SEM from three independent experiments. The p-value < 0.05 (*) was significant. (C, D and E) Huh-7.5 cells at 1.8 × 10^5^ cells in a twenty four-well plate were transfected with 24 pmol of either control siRNA or YAP1 siRNA. At 24 h after sinRNA-transfection, the cells were infected with HCV J6/JFH1 at an MOI of 2, cultured and harvested at the indicated time points. Total RNA was extracted, and YAP1, CYR61 and CTGF mRNA levels were quantified by RT-qPCR, normalized to GAPDH mRNA levels. The mRNA levels at day 2 in the mock-infected group was arbitrarily expressed as 1.0. Data are expressed as means ± SEM from three independent experiments. The p-value < 0.05 (*) was significant.

To determine whether HCV-induced upregulation of CYR61 and CTGF depends specifically on YAP1 nuclear translocation, we used siRNA to knock down YAP1 mRNA in HCV-infected Huh-7.5 cells. YAP1 mRNA levels were successfully reduced by approximately 80% (Fig. 4C). The mRNA levels of CYR61 and CTGF in HCV-infected cells were significantly lower than in siYAP1-transfected HCV-infected cells (Fig. 4D and 4E). These results suggest that HCV promotes the upregulation of CYR61 and CTGF via YAP1 nuclear translocation. CYR61 and CTGF are known to play roles in tissue remodeling and cell proliferation, suggesting that the HCV-induced upregulation of CYR61 and CTGF genes contributes to HCV-related pathogenesis.

### CYR61 and CTGF expression levels are elevated in patients with chronic HCV infection

To further validate the HCV-induced upregulation of CYR61 and CTGF in a clinical setting, we analyzed publicly available transcriptomic data. Both CYR61 and CTGF mRNAs were significantly upregulated in HCV-infected individuals compared to healthy controls (Fig. 5A and 5B), confirming upregulation of CYR61 and CTGF mRNAs in HCV-infected patients.

**Figure 5.**
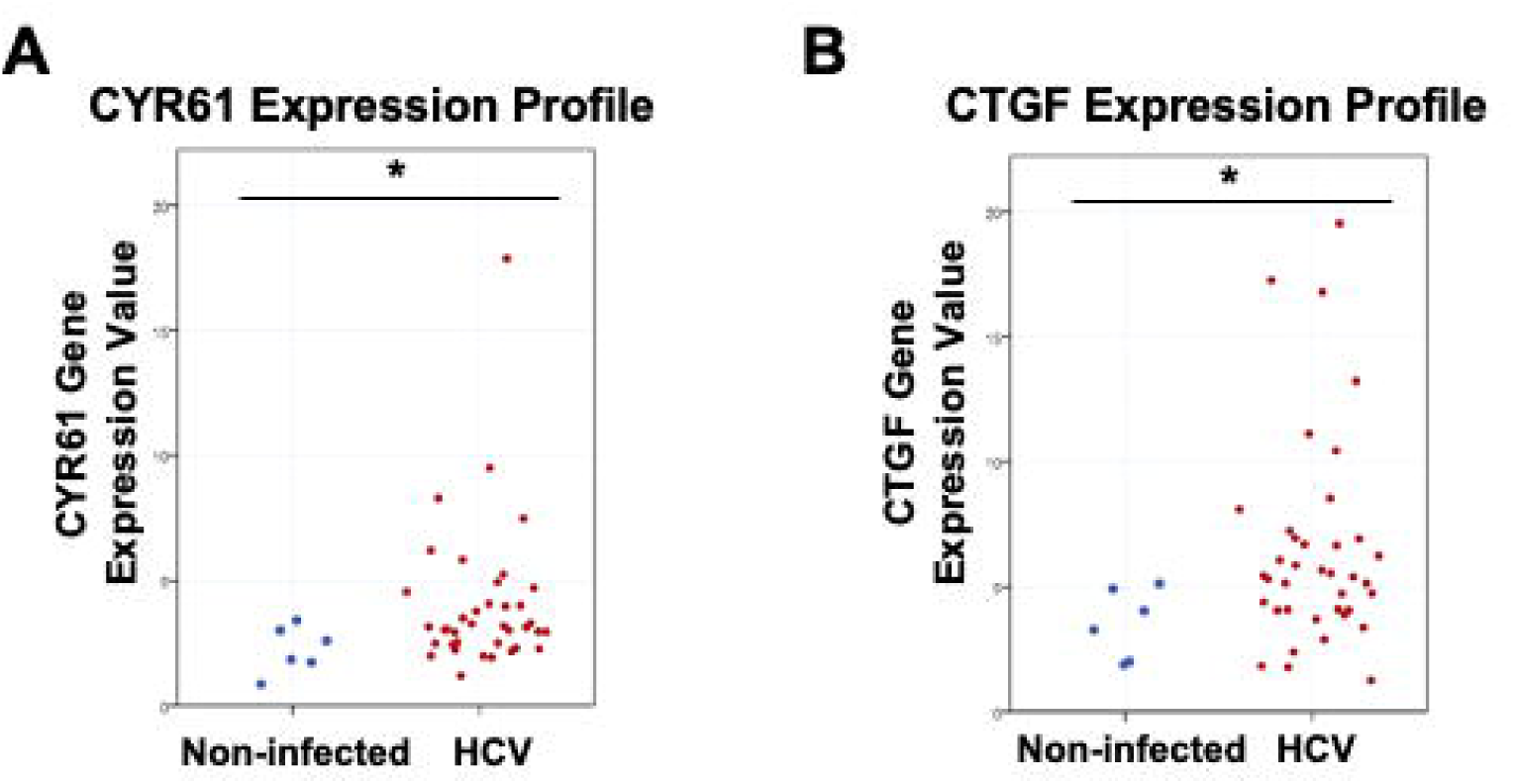
CYR61 and CTGF mRNA levels are increased in patients with chronic HCV infection. (A, B) CYR61 and CTGF mRNA levels in the liver tissue of patients with chronic HCV infection (n=38) compared to healthy individuals (n=6) (GEO accession number GSE 84346). Data are presented as the mean ± SD. The p-value < 0.05 (*) was significant.

In summary, we propose a model shown in Fig. 6. Under normal conditions, the Hippo pathway remains active when LATS1 phosphorylates YAP1, and phosphorylated YAP1 localizes in the cytoplasm. In HCV infection, HCV activates the ROS/JNK/Itch pathway (18). Active Itch induces the polyubiquitylation and proteasomal degradation of LATS1. The degradation of LATS1 inhibits YAP1 phosphorylation and promotes YAP1 nuclear translocation, thereby upregulating transcription of the YAP1 target genes, CYR61 and CTGF. HCV-induced suppression of the Hippo pathway plays a role in HCV pathogenesis.

**Figure 6.**
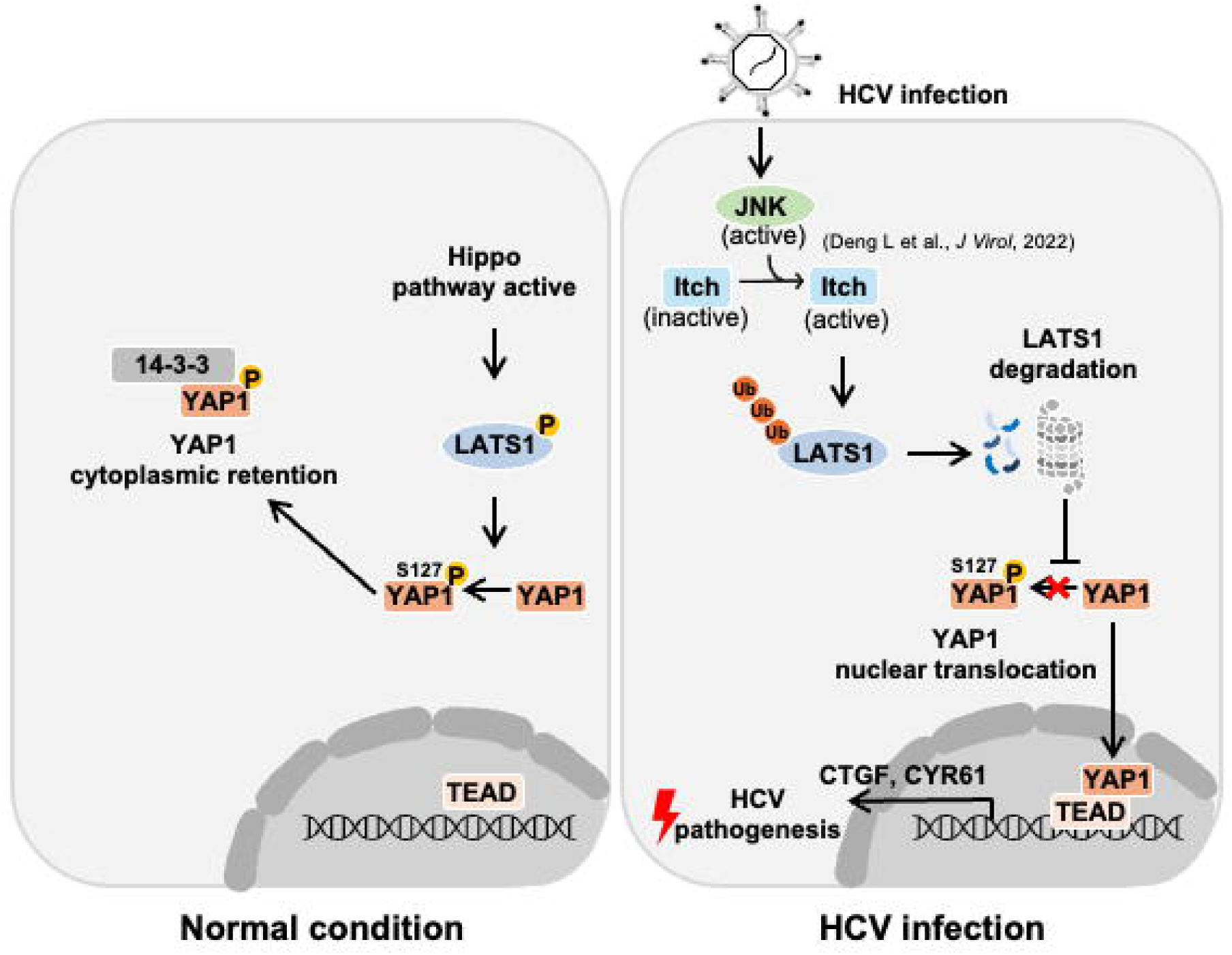
A proposed model of HCV-induced inactivation of the Hippo pathway and HCV-related pathogenesis. Under normal condition, the Hippo pathway remains active. LATS1 phosphorylates YAP1, thereby retaining YAP1 in the cytoplasm and preventing its nuclear entry. HCV infection activates the ROS/JNK/Itch signaling pathway, which leads to the phosphorylation and enhanced activity of Itch ubiquitin ligase. Activated Itch promotes the ubiquitin-dependent proteasomal degradation of LATS1 protein, leading to the Hippo pathway inactivation. As a result, YAP1 translocates from the cytoplasm to the nucleus and upregulates the transcription of CYR61 and CTGF genes, contributing to HCV-related pathogenesis.

## DISCUSSION

In this study, we found that HCV infection induced the degradation of LATS1 protein, a serine/threonine kinase essential for regulating the tumor-suppressive Hippo pathway (Fig. 1). We demonstrated that HCV infection inactivated the Hippo pathway and facilitated YAP1 nuclear translocation, and that HCV infection induced the upregulation of canonical YAP1 target genes, CYR61 and CTGF (Figs. 3-5).

Both CYR61 and CTGF are crucial for hepatocellular carcinoma (HCC) progression, exhibiting higher expression in HCC tissues than in normal liver tissues (27). CYR61 and CTGF act as adhesive substrates that stimulate fibroblast signaling, enhancing angiogenesis and wound healing (28). Notably, elevated CTGF levels in human HCCs have also been linked to poor clinical outcomes (29). We obtained evidence suggesting that HCV infection inactivates the Hippo pathway, leading to the upregulation of CYR61 and CTGF expression, thereby contributing to HCV pathogenesis.

The NEDD4 subfamily of HECT-type E3 ubiquitin ligases is characterized by multiple WW domains that specifically interact with the PPxY domain of LATS1. Notably, certain members of this subfamily have been identified as key regulators of LATS1 protein stability (19, 20, 30, 31). We hypothesized that the NEDD4 subfamily, such as Itch or WWP1, is involved in HCV-induced polyubiquitylation and proteasomal degradation of LATS1. We previously showed that HCV activates the ROS/JNK signaling pathway, which activates Itch and facilitates polyubiquitylation of VPS4A, enhancing HCV release (18). Our results suggest that Itch, but not WWP1, plays a crucial role in HCV-induced degradation of LATS1 protein (Fig. 2). Both Itch and WWP1 contributed to LATS1 polyubiquitylation in mock-infected cells. Importantly, siRNA-mediated knockdown of Itch restored LATS1 protein in HCV-infected cells. In contrast, siRNA-mediated knockdown of WWP1 further reduced LATS1 protein levels. These findings suggest that Itch predominantly plays a role in regulation of LATS1 protein during HCV infection, possibly because HCV activates Itch activity. WWP1 depletion may trigger compensatory activation of Itch, leading to enhanced LATS1 degradation rather than recovery.

Notably, condition-specific regulatory shifts have been observed in some proteins. For example, the tumor suppressor p53 is primarily degraded by Murine Double Minute 2 (Mdm2) under normal conditions, whereas in HPV16 E6-positive cancer cells, p53 degradation occurs entirely via the E6-AP pathway instead of Mdm2 (32). These observations raise the possibility that HCV infection predominantly facilitates Itch-mediated ubiquitylation and degradation of LATS1 via as-yet-undetermined mechanisms. Further investigation is required to elucidate the details of these regulatory mechanisms.

In this study, we demonstrated evidence suggesting that HCV infection promotes the ubiquitin-dependent degradation of LATS1 protein. This degradation contributes to the inactivation of the Hippo pathway, resulting in nuclear translocation of YAP1 and subsequent upregulation of the YAP1 target genes CYR61 and CTGF genes. To our knowledge, this is the first study to demonstrate that HCV-induced degradation of LATS1 leads to Hippo pathway inactivation, thereby upregulating CYR61 and CTGF genes. These insights may contribute to our understanding of the molecular mechanisms underlying HCV pathogenesis, highlighting potential targets for therapeutic intervention.

## AUTHOR STATEMENTS

### Author contributions

C.M. and I.S. conceived and designed the experiments. S.MA and C.M. carried out most of the experiments. ZX., F.P., C.M., L.D., and T.A. assisted the constructions and the data analysis. D.N.M.A. performed bioinformatics analysis of public gene expression data. S.MA., C.M, and I.S. wrote the manuscript.

### Conflicts of interest

The authors declare no conflict of interest.

## Funding information

This research was supported by grants for Basic and Clinical Research on Hepatitis from the Japan Agency for Medical Research and Development (AMED) grant No. 23fk0210090s1203 and 20fk0210040s0703, a KAKENHI grant No. 20K07514 and 22K15470, and the Program for the Nurturing of Next Generation Leaders Guiding Medical Innovation in Asia, funded by the Ministry of Education, Culture, Sports, Science, and Technology (MEXT) of Japan.

## Acknowledgements

We sincerely appreciate Dr. C. M Rice (The Rockefeller University, New York) for generously providing the Huh-7.5 cells and pFL-J6/JFH1 plasmid. We also thank Y. Kozaki for her valuable secretarial assistance.

## Notes

### Competing Interest Statement

The authors have declared no competing interest.

### Summary of Updates

Update Fig. 1C, 1D, 5A, and 5B as new data and modified the manuscript.

